## Supplemental Tables 1 to 3 for "Highly efficient CRISPR-Cas9-mediated gene knockout in primary human B cells for functional genetic studies of Epstein-Barr virus infection"

**Supplementary Table 1. List of Primer pairs**

| No | Gene | Strand | Nucleotide sequence | Purpose |
|---|---|---|---|---|
| 1 | CD46 | forward | CTGTACTACCTGCTGCCAGACC | Gel electrophoresis |
| 2 | CD46 | reverse | ATAACAGGCGTCATCTGAGACAGG | Gel electrophoresis |
| 3 | CD46 | forward | ACACTCTTTCCCTACACGACGctcttccgatctCTGTACTACCTGCTGCCAGACC | Miseq-single end seq |
| 4 | CD46 | reverse | TGACTGGAGTTCAGACGTGTGctcttccgatctATAACAGGCGTCATCTGAGACAGG | Miseq-single end seq |
| 5 | CD46 | forward | ACACTCTTTCCCTACACGACGctcttccgatctGACCACAGTCCATGGCTGATG | Miseq-paired end seq |
| 6 | CD46 | reverse | TGACTGGAGTTCAGACGTGTGctcttccgatctCATCACCGTAGTGGAATATGTACCC | Miseq-paired end seq |
| 7 | CDKN2A | forward | ACACTCTTTCCCTACACGACGctcttccgatctCGTCCTCCAGAGTCGCCC | Miseq-single end seq |
| 8 | CDKN2A | reverse | TGACTGGAGTTCAGACGTGTGctcttccgatctCTGCGGAGAGGGGGAGAG | Miseq-single end seq |

**Supplementary Table 2. crRNA sequences**

| Target gene | IDT design ID | Naming in the text | Nucleotide sequence | PAM sequence | Purpose |
|---|---|---|---|---|---|
| CD46 | Hs.Cas9.CD46.1.AA | CD46-gRNA 1 | TCGTTCACCAATCTCATAGT | AGG | CRISRP-Cas9 technology |
| CD46 | Hs.Cas9.CD46.1.AD | CD46-gRNA 2 | TTTGTGATCGGAATCATACA | TGG | CRISRP-Cas9 technology |
| CDKN2A | Hs.Cas9.CDKN2A.1.AA | CDKN2A-gRNA | CCCAACGCACCGAATAGTTA | CGG | CRISRP-Cas9 technology |

**Supplementary Table 3. Results of differential gene expression analysis of genes obtained by 4sU-RNA sequencing.**

| Number of genes | Gene name | padj |
|---|---|---|
| 1 | H3F3B | 0.0495853586509852 |
| 2 | EXOC6B | 0.0495722573540268 |
| 3 | FAM26F | 0.0494844708609299 |
| 4 | POLE3 | 0.0493864271426857 |
| 5 | ANKRD36BP2 | 0.0483248951400163 |
| 6 | FAM100B | 0.0483248951400163 |
| 7 | MRPS22 | 0.0481390340172488 |
| 8 | CACNA1A | 0.0480332231648256 |
| 9 | GPS2 | 0.0480332231648256 |
| 10 | SLC25A5 | 0.0480332231648256 |
| 11 | SLC7A5P2 | 0.0480332231648256 |
| 12 | TCL1A | 0.0480332231648256 |
| 13 | TPM4 | 0.0480332231648256 |
| 14 | CDKN2B | 0.047842682071218 |
| 15 | FBXL20 | 0.047842682071218 |
| 16 | MARCKSL1 | 0.0462407346491044 |
| 17 | PGAM1 | 0.046185347179266 |
| 18 | MYD88 | 0.0453305392198682 |
| 19 | RAPGEF5 | 0.0453305392198682 |
| 20 | SEMA4A | 0.0453305392198682 |
| 21 | BIRC5 | 0.0434579232466015 |
| 22 | LMO2 | 0.0429064262148305 |
| 23 | NLRP4 | 0.0429064262148305 |
| 24 | RBM7 | 0.0429064262148305 |
| 25 | KIF2C | 0.0416305651785589 |
| 26 | SYTL1 | 0.041078218420487 |
| 27 | RANBP1 | 0.0410228312489385 |
| 28 | GOLGA8A | 0.0392704675208476 |
| 29 | NCF4 | 0.0392704675208476 |
| 30 | RPL39 | 0.0392704675208476 |
| 31 | NAB2 | 0.0386714438695081 |
| 32 | HIST1H2AM | 0.0381753838397577 |
| 33 | HIST1H4B | 0.0381753838397577 |
| 34 | SNRPF | 0.0381753838397577 |
| 35 | ANTXR2 | 0.0378382503423391 |
| 36 | ATPIF1 | 0.0377662812363794 |
| 37 | COX17 | 0.0374978448337278 |
| 38 | LOC285758 | 0.0369456690557389 |
| 39 | EID1 | 0.0367800094849309 |
| 40 | INADL | 0.0367800094849309 |
| 41 | LCP1 | 0.0367800094849309 |
| 42 | SNX29 | 0.0356407389380155 |
| 43 | CD72 | 0.0352698806508784 |
| 44 | N4BP2 | 0.0352698806508784 |
| 45 | NUSAP1 | 0.0352698806508784 |
| 46 | FBXO5 | 0.0352233609586013 |
| 47 | TUBA1B | 0.0352233609586013 |
| 48 | IFITM2 | 0.0350946157929767 |
| 49 | MIF | 0.0344988064043864 |
| 50 | TMCO7 | 0.0340602417566875 |
| 51 | RPL14 | 0.0332747413675659 |
| 52 | AAK1 | 0.0329404123308122 |
| 53 | HIST1H2BH | 0.0328162212868041 |
| 54 | ING3 | 0.0320175269138185 |
| 55 | LOC100505812 | 0.0319385371903286 |
| 56 | CBR1 | 0.0316305645183963 |
| 57 | CHI3L2 | 0.0295101636278342 |
| 58 | LOC646329 | 0.0295101636278342 |
| 59 | RPL8 | 0.0292436989028342 |
| 60 | DCK | 0.0291961519789457 |
| 61 | MAD2L1 | 0.0291101012463462 |
| 62 | HMGB2 | 0.0280460594795761 |
| 63 | SUMO3 | 0.0274356798140161 |
| 64 | C4orf46 | 0.0270576567173824 |
| 65 | LYSMD2 | 0.0270576567173824 |
| 66 | PSMA4 | 0.0270576567173824 |
| 67 | SMS | 0.0270576567173824 |
| 68 | SULF2 | 0.0270576567173824 |
| 69 | TPI1 | 0.0270576567173824 |
| 70 | HIST1H2BN | 0.0256002811571811 |
| 71 | KIAA0040 | 0.0256002811571811 |
| 72 | CAP1 | 0.024298401624602 |
| 73 | MAP2K6 | 0.024298401624602 |
| 74 | HDAC5 | 0.0242919975648105 |
| 75 | HIST1H2BL | 0.0242919975648105 |
| 76 | HIST1H4D | 0.0237866514479686 |
| 77 | CCNB2 | 0.0228375384436799 |
| 78 | HIST2H3D | 0.0223153810170486 |
| 79 | ARPC5 | 0.0210455459049929 |
| 80 | C14orf2 | 0.0210455459049929 |
| 81 | IDH3A | 0.0210455459049929 |
| 82 | HIST1H3G | 0.0210362715710858 |
| 83 | LSM5 | 0.0210362715710858 |
| 84 | PSME2 | 0.0210362715710858 |
| 85 | TCEAL8 | 0.0210362715710858 |
| 86 | CD79A | 0.0209769657049205 |
| 87 | CCL4 | 0.02090268482058 |
| 88 | COX6A1 | 0.0205500444710553 |
| 89 | SNRPD1 | 0.0205500444710553 |
| 90 | ACTG1 | 0.0205249820474698 |
| 91 | IFIT5 | 0.0205249820474698 |
| 92 | ZFYVE9 | 0.0205249820474698 |
| 93 | CKS2 | 0.0199227331765973 |
| 94 | DDB2 | 0.0197229079052022 |
| 95 | C1QBP | 0.0197147028626663 |
| 96 | NACA | 0.019300044395372 |
| 97 | GAPDH | 0.0191457748670181 |
| 98 | MT2A | 0.018960957500115 |
| 99 | RPS20 | 0.018960957500115 |
| 100 | LRRC32 | 0.0187982660002832 |
| 101 | TNRC6C | 0.0186290558583279 |
| 102 | HIST1H2BI | 0.018470751571979 |
| 103 | RPL22 | 0.0180803761047385 |
| 104 | TMOD2 | 0.0176438826104996 |
| 105 | HIST1H4C | 0.0169800531944221 |
| 106 | FAM18B2 | 0.014424040526018 |
| 107 | BCL11A | 0.0137600910666385 |
| 108 | HIST1H3H | 0.0130640459490103 |
| 109 | CCL3 | 0.0126913996635382 |
| 110 | CYP1B1 | 0.0126913996635382 |
| 111 | IRF8 | 0.0126913996635382 |
| 112 | IRS2 | 0.0126913996635382 |
| 113 | RPL35 | 0.0126913996635382 |
| 114 | TBCC | 0.0126913996635382 |
| 115 | CCPG1 | 0.0124899488026773 |
| 116 | CFL1 | 0.0124899488026773 |
| 117 | ELL3 | 0.0124703472322331 |
| 118 | HIST2H2AC | 0.0124703472322331 |
| 119 | HIST1H2BD | 0.0124454815594565 |
| 120 | RPL19 | 0.0113408091931427 |
| 121 | UQCR10 | 0.0113408091931427 |
| 122 | HIST1H2BJ | 0.0111908502217577 |
| 123 | LY6E | 0.0101240989421758 |
| 124 | SMN1 | 0.0101240989421758 |
| 125 | HIST1H1E | 0.0100107296285801 |
| 126 | HIST1H2AL | 0.0100107296285801 |
| 127 | RAC2 | 0.00985646928899781 |
| 128 | FLJ43663 | 0.00974323936413508 |
| 129 | BCL2A1 | 0.00937831974247774 |
| 130 | LSM6 | 0.00937831974247774 |
| 131 | CD74 | 0.00590983018253672 |
| 132 | PVT1 | 0.00590983018253672 |
| 133 | HIST1H2AJ | 0.00567283270398546 |
| 134 | HIST1H2BF | 0.00567283270398546 |
| 135 | MRPS18B | 0.00567283270398546 |
| 136 | VAV3 | 0.00562233294490733 |
| 137 | PFN1 | 0.00539459191246031 |
| 138 | RPL23A | 0.00511094895956291 |
| 139 | RASSF6 | 0.00500080480548445 |
| 140 | NFKBIE | 0.00350045257636285 |
| 141 | TREX1 | 0.00350045257636285 |
| 142 | PDE4B | 0.00343579054199575 |
| 143 | PRDM8 | 0.00335984834029918 |
| 144 | CCNB1 | 0.00334160950091299 |
| 145 | BACH2 | 0.00315723082229448 |
| 146 | IFIT3 | 0.00284106233353294 |
| 147 | COX6C | 0.00282678622347304 |
| 148 | HIST1H2AE | 0.00280890906530428 |
| 149 | HIST1H2BK | 0.00280890906530428 |
| 150 | TMSB4X | 0.00280890906530428 |
| 151 | CKAP2L | 0.0025586265986473 |
| 152 | ASTN2 | 0.00195020467131566 |
| 153 | DUSP2 | 0.00195020467131566 |
| 154 | EGR3 | 0.00195020467131566 |
| 155 | PNPLA7 | 0.00195020467131566 |
| 156 | SAT1 | 0.00195020467131566 |
| 157 | EID3 | 0.00188555341578105 |
| 158 | PSME1 | 0.00188555341578105 |
| 159 | ACTB | 0.00139109987757177 |
| 160 | IGLL5 | 0.00127388444193456 |
| 161 | HIST1H2BB | 0.00107127022610895 |
| 162 | PTMA | 0.00107127022610895 |
| 163 | SSTR2 | 0.00107127022610895 |
| 164 | STARD9 | 0.00107127022610895 |
| 165 | ZDHHC14 | 0.00104031773750262 |
| 166 | SBF2 | 0.000952973362878684 |
| 167 | TMEM163 | 0.000952973362878684 |
| 168 | COX7C | 0.000945086067370795 |
| 169 | HIST1H4F | 0.000945086067370795 |
| 170 | CCDC88B | 0.000616176839499103 |
| 171 | IFI30 | 0.000616176839499103 |
| 172 | RPL36AL | 0.000575857264211619 |
| 173 | SERPINA9 | 0.000551511264784517 |
| 174 | HIST2H2AB | 0.000389510668245257 |
| 175 | HIST1H2AG | 4,44E+09 |
| 176 | HIST1H2BC | 4,27E+09 |
| 177 | MYL12A | 6,07E+07 |
| 178 | LTA | 5,64E+07 |
| 179 | HIST2H2BF | 3,37E+08 |
| 180 | DNAH7 | 2,63E+08 |
| 181 | LINC00152 | 3,86E+07 |
| 182 | PTPN6 | 3,62E+06 |
