## Supplemental Figures S1 to S4 for "Highly efficient CRISPR-Cas9-mediated gene knockout in primary human B cells for functional genetic studies of Epstein-Barr virus infection"

A

MiSeq sequencing and data analysis of CD46-Cas9 cells and controls

Outknocker web tool

CD46 Cas9 cells:
Knockout efficiency: 84.2 %

WT cells:
Knockout efficiency: 0.1 %

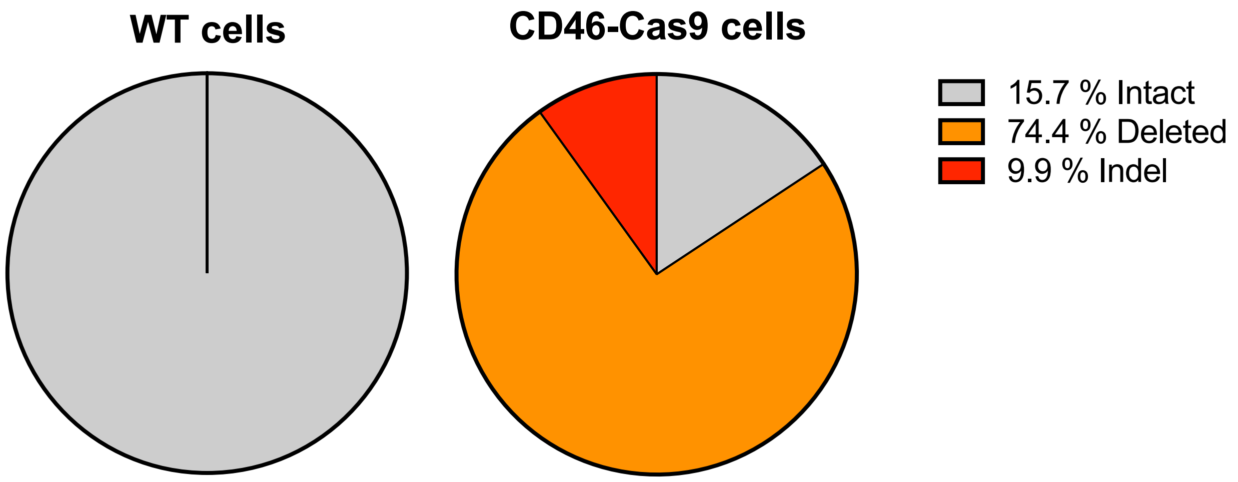


| | | |
|---|---|---|
| | PAM | |
| REFERENCE | CTATGGAGCTCATTGGTAAACCAAAACCCTACTATGAGATTGGTGAACGAGTAGATTATAAGTGTAAAAAAGGATACTTC | |
| Intact | | **15.71%** |
| no indel | CTATGGAGCTCATTGGTAAACCAAAACCCTACTATGAGATTGGTGAACGAGTAGATTATAAGTGTAAAAAAGGATACTTC | 919 reads |
| Deleted | | **74.35%** |
| 90nt deletion | CTATGGAGCTCATTGGTAAACCAAAACCCTACTAT | 274 reads |
| 91nt deletion | CTATGGAGCTCATTGGTAAACCAAAACCCTACT | 2995 reads |
| 92nt deletion | CTATGGAGCTCATTGGTAAACCAAAACCCTACTA | 1081 reads |
| Indel | | **9.94%** |
| 1nt deletion | CTATGGAGCTCATTGGTAAACCAAAACCCTACT_TGAGATTGGTGAACGAGTAGATTATAAGTGTAAAAAAGGATACTTC | 316 reads |
| 2nt deletion | CTATGGAGCTCATTGGTAAACCAAAACCCTACT__GAGATTGGTGAACGAGTAGATTATAAGTGTAAAAAAGGATACTTC | 265 reads |

B

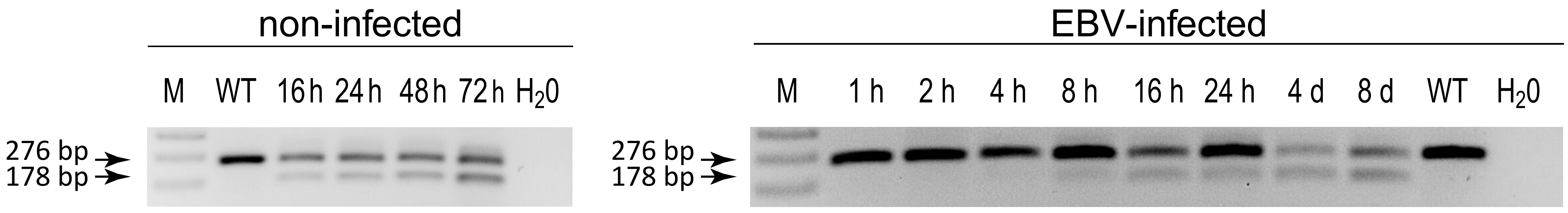


Supplementary Figure S1

### MiSeq sequencing and data analysis of p16 KO cells and controls

Outknocker web tool

p16 KO cells:
Knockout efficiency: 74.6 %

WT cells:
Knockout efficiency: 0.1 %

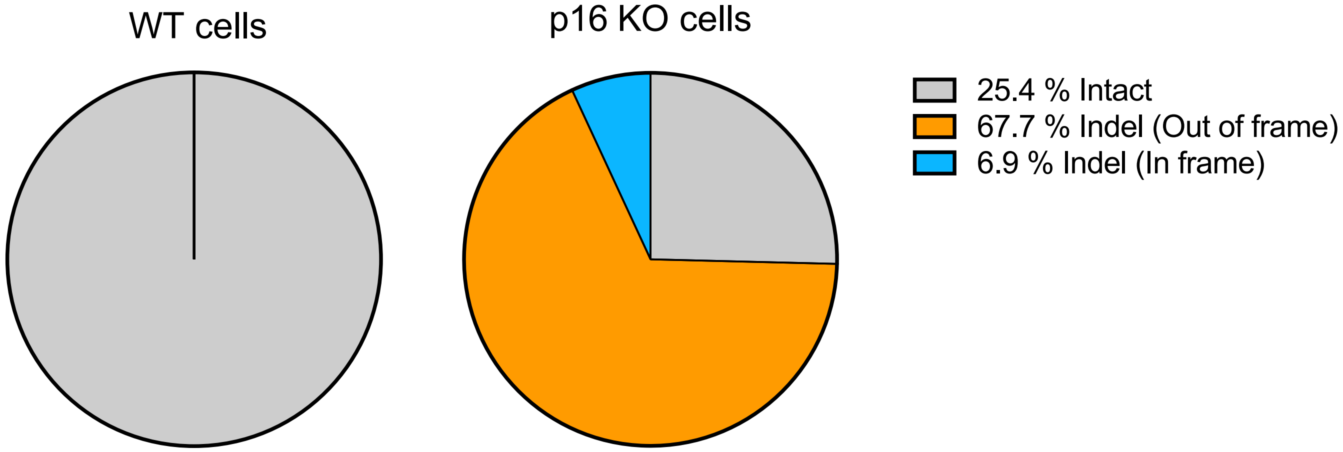


| | | |
|---|---|---|
| | PAM | |
| REFERENCE | CCTCTACCCACCTGGATCGGCCTCCGACCGTAACTATTCGGTGCGTTGGGCAGCGCCCCGCCTCCAGCAGCGCCCGCAC | |
| Intact | | **25.4 %** |
| no indel | CCTCTACCCACCTGGATCGGCCTCCGACCGTAACTATTCGGTGCGTTGGGCAGCGCCCCGCCTCCAGCAGCGCCCGCAC | 1145 reads |
| Indel (Out of frame) | | **67.7 %** |
| 1nt deletion | CCTCTACCCACCTGGATCGGCCTCCGACCGTAACTA_TCGGTGCGTTGGGCAGCGCCCCGCCTCCAGCAGCGCCCGCAC | 702 reads |
| | CCTCTACCCACCTGGATCGGCCTCCGACCGTA_CTATTCGGTGCGTTGGGCAGCGCCCCGCCTCCAGCAGCGCCCGCAC | |
| 1nt insertion | CCTCTACCCACCTGGATCGGCCTCCGACCGTAAACTATTCGGTGCGTTGGGCAGCGCCCCGCCTCCAGCAGCGCCCGCAC | |
| 2nt deletion | CCTCTACCCACCTGGATCGGCCTCCGACCGTAA__ATTCGGTGCGTTGGGCAGCGCCCCGCCTCCAGCAGCGCCCGCAC | |
| 4nt deletion | CCTCTACCCACCTGGATCGGCCTCCGACCGTAA____CGGTGCGTTGGGCAGCGCCCCGCCTCCAGCAGCGCCCGCAC | |
| | CCTCTACCCACCTGGATCGGCCTCCGACCGTA____TTCGGTGCGTTGGGCAGCGCCCCGCCTCCAGCAGCGCCCGCAC | |
| 5nt deletion | CCTCTACCCACCTGGATCGGCCTCCGACCGTAAC_____GGTGCGTTGGGCAGCGCCCCGCCTCCAGCAGCGCCCGCAC | |
| Indel (In frame) | | **6.9 %** |
| 3nt deletion | CCTCTACCCACCTGGATCGGCCTCCGACCGTAA___TTCGGTGCGTTGGGCAGCGCCCCGCCTCCAGCAGCGCCCGCAC | 316 reads |

Supplementary Figure S2

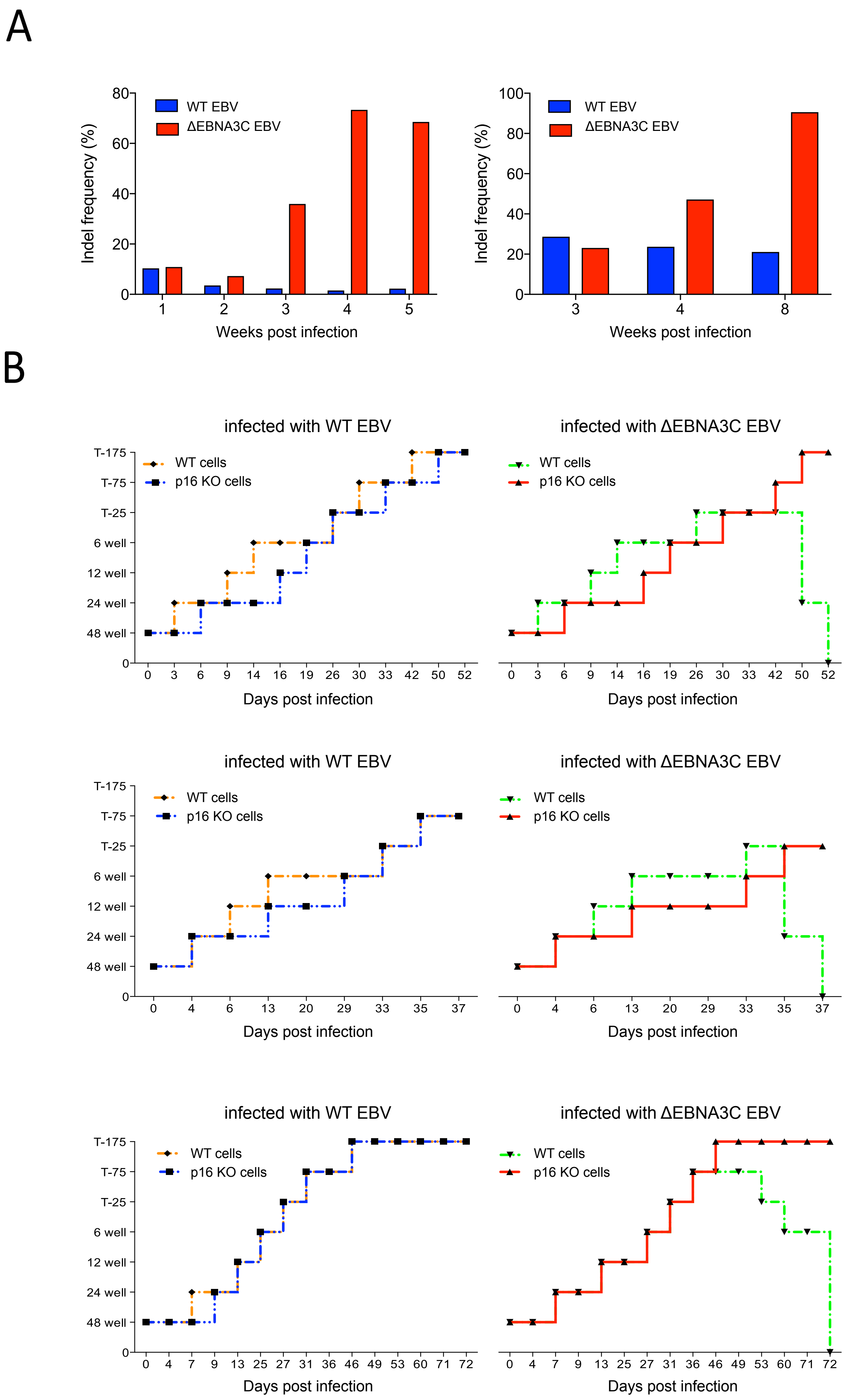


Supplementary Figure S3

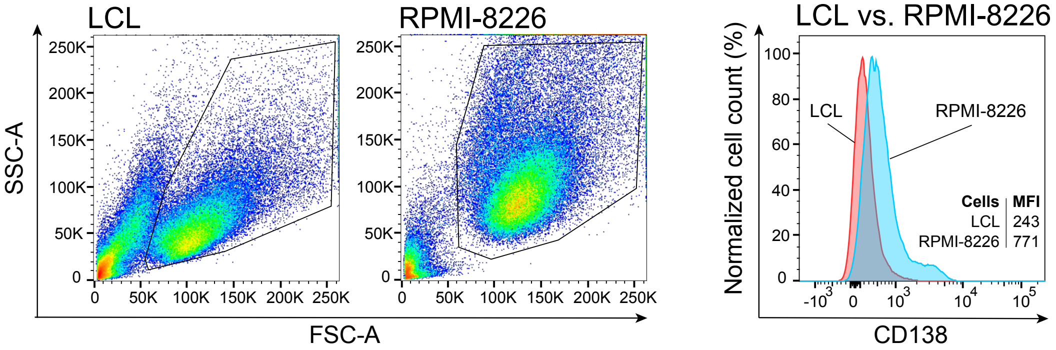


### Supplementary Figure S4
